# Prioritizing transcriptomic and epigenomic experiments by using an optimization strategy that leverages imputed data

**DOI:** 10.1101/708107

**Authors:** Jacob Schreiber, Jeffrey Bilmes, William Stafford Noble

## Abstract

Successful science often involves not only performing experiments well, but also choosing well among many possible experiments. In a hypothesis generation setting, choosing an experiment well means choosing an experiment whose results are interesting or novel. In this work, we formalize this selection procedure in the context of genomics and epigenomics data generation. Specifically, we consider the task faced by a scientific consortium such as the National Institutes of Health ENCODE Consortium, whose goal is to characterize all of the functional elements in the human genome. Given a list of possible cell types or tissue types (“biosamples”) and a list of possible high throughput sequencing assays, we ask “Which experiments should ENCODE perform next?” We demonstrate how to represent this task as an optimization problem, where the goal is to maximize the information gained in each successive experiment. Compared with previous work that has addressed a similar problem, our approach has the advantage that it can use imputed data to tailor the selected list of experiments based on data collected previously by the consortium. We demonstrate the utility of our proposed method in simulations, and we provide a general software framework, named Kiwano, for selecting genomic and epigenomic experiments.

## 1 Introduction

Experimental characterization of the genomic and epigenomic landscape of a human cell line or tissue (“biosample”) is expensive but can potentially yield valuable insights into the molecular basis for development and disease. Fully measuring the epigenome involves, in principle, assaying chromatin accessibility, transcription, dozens of histone modifications, and the binding of over a thousand DNA-binding proteins. Even after accounting for the decreasing cost of high-throughput sequencing, such an exhaustive analysis is expensive, and systematically applying such techniques to diverse cell types and cell states is simply infeasible. Essentially, we cannot afford to fill in an experimental data matrix in which rows correspond to types of assays and columns correspond to biosamples.

Several approaches have been proposed to address this challenge. Some scientific consortia, such as GTEx and ENTEX, aim to completely fill in a submatrix of selected assays and selected biosamples. In contrast, other consortia, such as the Roadmap Epigenomics Mapping Consortium [1] and ENCODE [2], adopted a roughly “L”-shaped strategy, in which consortium members focused on carrying out many assays in a small set of high-priority biosamples, and some assays were carried out over a much larger set of biosamples. Recently, computational approaches have been proposed that rely on using machine learning models to impute the experiments that have not yet been performed [3, 4, 5, 6]. While the imputation strategy can relatively easily complete the entire matrix, a drawback is that the imputed data is potentially less trustworthy than actual experimental data.

In this work, we address a variant of the matrix completion problem. Specifically, we consider the scenario that we, as a field, find ourselves in currently, having performed many assays in many biosamples and trying to figure out which of the remaining assay/biosample combinations (“experiments”) we should perform next. In many cases, the choice of which experiments to perform is driven by intuitition and guesswork. We hypothesize that a data-driven approach to this problem can increase the rate of scientific discovery.

Previous work by Wei et al. [7] has addressed a closely related problem. Wei et al. studied the problem of filling in a new row (or column) of the matrix. Say that we have decided to begin performing experiments on a new biosample, but we can only afford to carry out a fixed number *k* of assays. Then the question is, “Which set of *k* assays is likely to yield the most information?” Wei et al. answer this question by framing the task as an optimization problem, where we attempt to maximize a function *f* (·) that quantifies the joint quality of a given subset of size *k* relative to the full collection of possible assays.

An important feature of the method proposed by Wei et al. is that the approach is “cell-type agnostic,” in the sense that it yields a single set of suggested assays, irrespective of what biosample will be analyzed. This property arises because the set quality function *f* (·) measures the similarity of a given pair of assays by averaging across all cell types in which both assays have already been performed. Wei et al. explicitly consider the scenario in which a specified set of assays has already been performed in a given biosample, and the task is to select the next *k* assays to perform. However, even in this setting, the proposed approach is cell-type agnostic: the method yields the same answer for any biosample in which the specified set of assays has been performed.

In this work, we propose a method that can select experiments that span a diverse set of biosamples and assays jointly. The key idea is to apply the quality function *f* (·) to similarities calculated using imputed, rather than real, data. The resulting method, implemented in a software package called “Kiwano,” is far more flexible and powerful than the original method. Most importantly, rather than restricting our selection to a single row or column of the data matrix, using imputed data allows us to address the global question, “Among all possible experiments, which one should I do next?” Furthermore, even in the case where we want to select assays to perform in a given biosample, our imputation-based approach selects a set that is tailored to this particular biosample, by computing similarities between the imputed values for potential experiments and experiments that have already been performed in that biosample.

We validate Kiwano in several ways, using ENCODE data. First, we demonstrate via visualization that the imputation-based similarity matrix encodes meaningful biological relationships among assay types and biosamples. We then apply the optimization procedure to this similarity matrix and show that the resulting subset of experiments is representative of the full set, both qualitatively and through simulation experiments. We also illustrate how to apply the objective function used in our optimization to ascertain which biosamples are currently undercharacterized or which assays are underutilized. We have made a tool available at https://www.github.com/jmschrei/kiwano/ that can order experiments based on the pre-calculated similarity matrix we use here.

## 2 Results

### 2.1 Imputations cluster according to known biological patterns

Our approach for prioritizing experimental characterization relies on a similarity matrix that is calculated on imputed experiments. To produce this matrix, we first generated imputations of epigenomic and transcriptomic experiments using a recently developed imputation approached based on deep tensor factorization, named Avocado. These imputations span 400 human biosamples and 77 assays of biological activity for a total of 30,800 imputed tracks. After acquiring these imputations, we calculated the squared Pearson correlation between all pairs of imputed experiments for use as a similarity measure, resulting in a 30,800 by 30,800 matrix.

After calculating the similarity matrix, we investigated whether the similarity matrix was able to capture high level biological trends that would be crucial for prioritization. We began by visually inspecting a two-dimensional UMAP projection of the similarity matrix down to two dimensions (Figure 1a). The clearest trend in this projection is a separation of experiments based on a broad categorization of the type of activity measured by the assay. We observed that one cluster contained mostly protein binding experiments, one contained mostly histone modification experiments, and several neighboring clusters were composed exclusively of transcription-measuring experiments. Initially, one might expect that experiments in the same biosample where the assays measure the same underlying phenomena might cluster together. However, we observed that in some cases a pair of experiments may exhibit low correlation when the shape of their signals along the genome differ, even when the assays used in the experiments both measure the same underlying biological activity. For example, the histone modification H3K36me3 is known to be associated with transcription but generally forms broad peaks across the entire gene body, whereas assays such as CAGE or RAMPAGE form punctate peaks.

**Figure 1:**
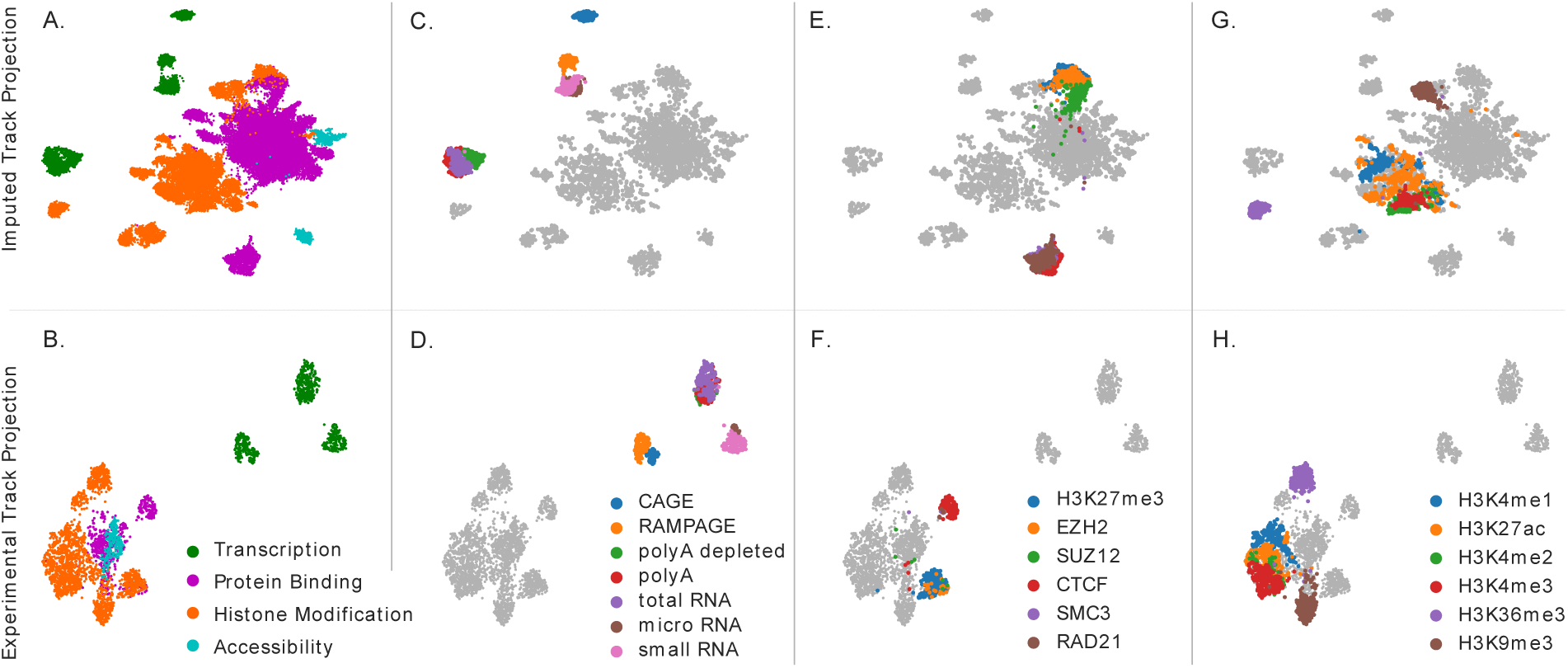
A projection of imputed and experimental epigenomic tracks. Each panel shows a UMAP projection of 30,800 imputed experiments (top row) or of 3,150 tracks of primary data (bottom row). In each column, a different set of experiments is highlighted based on their biological activity. (A/B) Experiments are highlighted based on broad categorization of the assayed activity. (C/D) Transcription measuring experiments are colored according to different types of assays. (E/F) Experiments are highlighted that measure H3K27me3 and two polycomb subunits, as well as CTCF and two cohesin subunits. (G/H) Experiments are highlighted showing several histone modifications that are enhancer-associated, such as H3K4me1 (blue) and H3K27ac (orange), promoter-associated such as H3K4me2 (green) and H3K4me3 (red), transcription-associated such as H3K36me3 (purple), or broadly repressive such as H3K9me3 (brown).

In order to confirm that the separation according to assay categorization was not an artifact of the imputation process, we used the same process to calculate a similarity matrix and subsequent UMAP projection for the 3,150 tracks of the experimental (or “primary”) data (Figure 1b). The major trends present in the projection of imputed data are consistent with those in the primary data. In particular, transcription experiments form distinct clusters, protein binding experiments are mostly distinct from histone modification ones, and chromatin accessibility experiments localize closer to protein binding experiments than to histone modification experiments. Note that although the figure may appear to show that accessibility experiments overlap with protein binding experiments, a closer examination reveals that the protein binding experiments mostly surround the accessibility experiments.

Next, we more closely examined four sets of assays that, a priori, we expected to show distinctive patterns. The first set of experiments were those that measured transcription. When we highlighted experiments by assay type, we observed CAGE and RAMPAGE experiments forming distinct cluters, micro- and small-RNA-seq experiments forming a third cluster, and polyA-, polyA-depleted-, and total-RNA-seq experiments forming a fourth (Figure 1c/d). The second and third sets of experiments involved triplets of assays whose activity are usually associated, specifically, with CTCF and the cohesin subunits, SMC3 and RAD21, as well as H3K27me3 and two polycomb subunits, EZH2 and SUZ12 (Figure 1e/f). In both cases we observe distinct clusters of experiments, which is particularly interesting for H3K27me3 and the polycomb subunits because one assay measures a histone modification and the other two measure protein binding. The fourth set of experiments focused on six well-studied histone modifications (Figure 1g/h). The clustering of these six marks coincide with the genomic element in which they are typically enriched. In particular, experiments measuring H3K36me3 and H3K9me3 form their own clusters, with the two assays respectively measuring activity enriched in gene bodies and constitutive heterochromatin. Further, the primary cluster of histone modification experiments exhibited a separation between the promoter-associated marks, H3K4me2 and H3K4me3, and the enhancer-associated marks, H3K4me1 and H3K27ac. We observed similar patterns across both the imputed and primary data for each of these four sets of assays. Taken together, these observations suggest that a similarity matrix derived from imputed experiments is successfully capturing important aspects of real biological activity.

### 2.2 Submodular selection of imputations flexibly prioritizes assays across cellular contexts

Having shown that the similarity matrix captures several high level trends in the data, we turn to the task of experimental prioritization. Our strategy for prioritizing experiments relies on submodular selection, which is a technique for reducing a set of elements to a minimally redundant subset through the optimization of a submodular function that captures the quality, or “representativeness,” of a given subset relative to the full set (see Methods for details). Submodular selection has been used previously to select genomics assays [7], to select representative sets of protein sequences [8], and to choose genomic loci for characterization by CRISPR-based screens [9]. Specifically, we optimize a “facility location” objective function, which operates on pairwise similarities between elements and so is well suited to leverage our similarity matrix (see Methods).

A critical property of submodular functions is that greedy optimization will yield a subset whose objective value is within 1 − *e*^− 1^ of the optimal subset, and that this is the best approximation one can make unless P=NP [10]. This greedy optimization algorithm iteratively selects the single element whose inclusion in the representative set leads to the largest gain in the objective function. Thus, when applied to our similarity matrix, the submodular selection procedure will yield an ordering over all experiments that attempts to minimize redundancy among those experiments that are selected early in the process.

In order to demonstrate that submodular selection results in a representative subset of assays, we applied it to our calculated similarity matrix. Visually, we observe that the first 50 selected experiments appear to cover the space well and include selections from many of the small clusters of experiments (Figure 2a, Supplementary Table 1). When we count the number of assays selected for each type of biological activity, we find that protein binding assays are the most commonly selected with 23 experiments, followed by histone modification assays with 19 experiments, transcription assays with 6 experiments, and, finally, accessibility assays with 2 experiments (Figure 2b). However, when we compare the number of selected experiments of each type to the number that one would expect by randomly selecting with replacement, we observe that protein binding experiments are underrepresented, whereas histone modification experiments are overrepresented. We note that the first 10 experiments are at the centers of large clusters of experiments and that the subsequent 40 experiments are selected from smaller clusters. This finding corresponds with the gain in the facility location objective score from each successive experiment significantly diminishing by the tenth experiment (Figure 2c).

**Figure 2:**
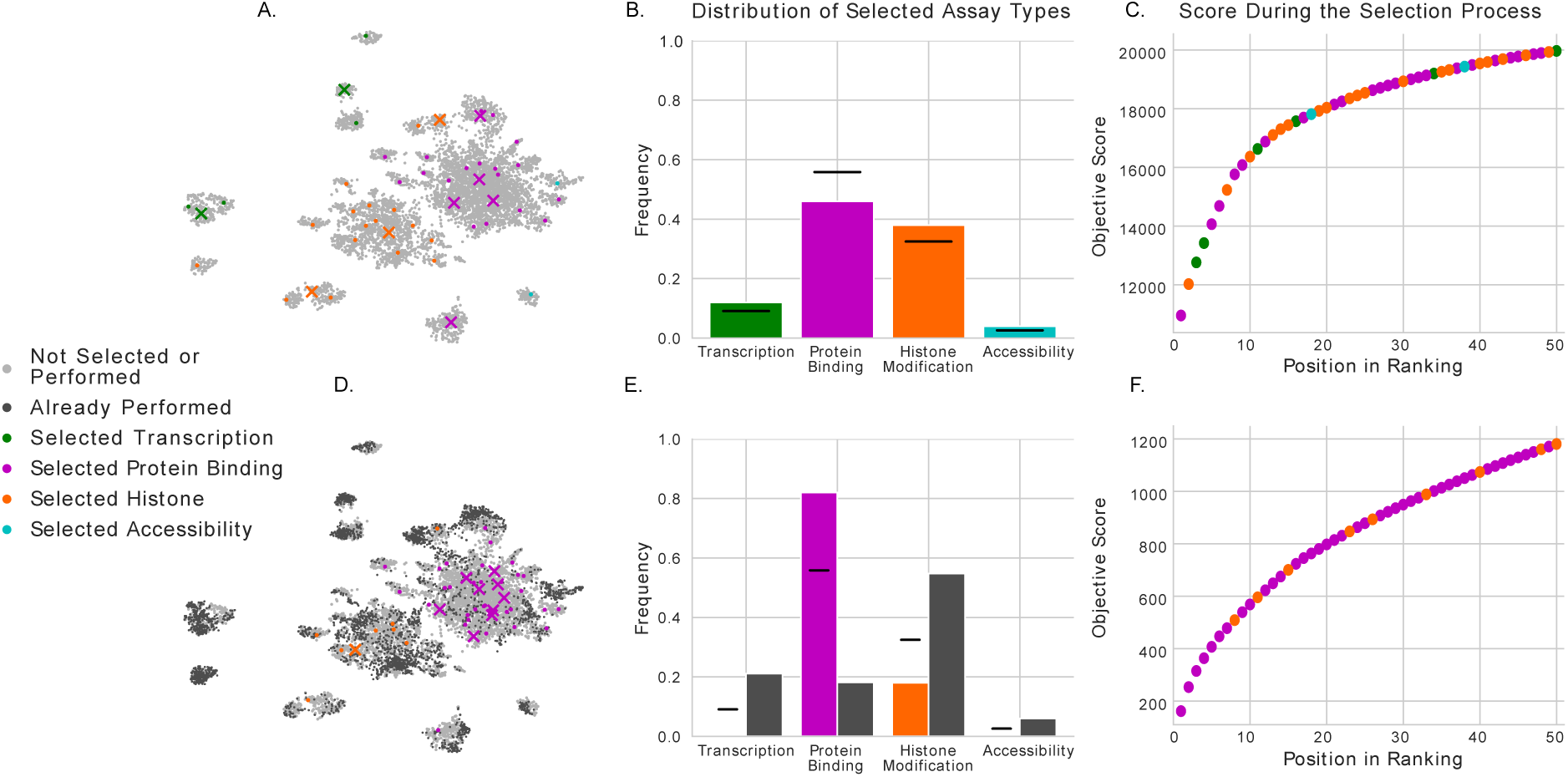
A selection of experiments before and after accounting for those that have already been performed. (A) The same projection of imputed experiments as shown in Figure 1a, where the first 50 experiments selected using Kiwano are colored by the type of activity that they measure. The first 10 experiments selected are marked using an X, and the remaining 40 are marked with a dot. (B) A bar chart showing the frequency that experiments of each type of activity are selected in the first 50 experiments. (C) The facility location objective score as the first 50 experiments are selected, with each point colored by the type of activity measured by that experiment. (D) The same as (A), but with the selection procedure initialized with the experiments that have already been performed, and with those experiments displayed in dark grey. (E) The same as (B), but with dark grey bars showing the frequency of experiments of each type that have already been performed. (F) The same as (C), but with the selection procedure initialized with the experiments that have already been performed.

A weakness in simply applying submodular selection to the full set of imputed experiments is that the procedure does not account for the thousands of epigenomic and transcriptomic experiments that have already been performed. Fortunately, there are two ways that one can account for these experiments. The first is to remove those experiments that have already been performed from the similarity matrix and perform selection on the remaining experiments. While this approach is simple, it does not account for the content of the experiments that have already been performed. For example, if transcription has already been measured in hundreds of biosamples, then it may be beneficial to focus experimental efforts on characterizing other types of biological activity. A second approach takes advantage of the fact that the selection process is greedy by initializing the set of selected experiments with those that have already been performed. This ensures that the selected experiments cover types of activity that are not already well characterized.

Accordingly, we proceeded with the second approach. We initialized a facility location function with the 3,150 experiments that had already been performed and ranked the remaining 27,650 experiments. We observed that the selected experiments lie primarily in areas of the UMAP projection that do not already have many experiments performed (Figure 2d, Supplementary Table 2). When we counted the number of selected experiments of each type, we found that the number of protein binding experiments increased from 23, when not accounting for the experiments that had already been performed, to 41, when accounting for them (Figure 2e). Correspondingly, the number of histone modification experiments decreased from 19 to 9. This change in coverage is likely because 1,726 experiments measuring histone modification have already been performed, whereas only 571 experiments measuring protein binding have been performed. Further, none of the first 50 selected experiments measure transcription or accessibility, likely because those forms of activity are already much better measured than protein binding. In this setting, the gain in the facility location objective function of each successive experiment is much lower, due in large part to the experiments that have already been performed (Figure 2f).

### 2.3 Selection on imputed experiments identify diversity in primary data

Our next step was to evaluate the quality of the selected experiments in a quantitative way. Following Wei et al; we reasoned that the signal contained in a representative subset of experiments would be well suited for reconstructing the signal in all experiments. We formulated the problem of quantitatively measuring how representative a subset is as a multi-task regression problem, with the input features being the signal from the selected subset of experiments and the outputs being the signal from the full set of experiments (see Methods). Importantly, to ensure that this validation measured how representative a subset is of the primary data, despite subset selection having been performed on the imputations, we used the primary data as both the input and target for this task.

We selected a subset of experiments in three ways. The first was through the submodular selection procedure described in Section 2.2, applied to the 3,150 imputed experiments for which primary data had already been collected. The second was by applying the submodular selection procedure to the 3,150 tracks of primary data themselves. Naturally, selecting subsets based on the primary data cannot be extended to experiments that have not yet been performed, and so the purpose of evaluating models trained using this subset is to measure the effect that the imputation process itself has on selecting a representative subset of experiments. The third was selecting subsets of the 3,150 performed experiments at random. This random process was repeated 20 times to obtain a distribution of scores.

We observe that the subsets of experiments selected using submodular selection consistently outperform those selected at random (Figure 3). Each comparison is statistically significant at a p-value threshold of 0.01 according to a one sample t-test. Further, for smaller subsets, applying submodular selection to the imputed tracks performs nearly as well as the panels selected on the primary data itself, showing that the distortion introduced by the imputation process is small. Interestingly, when the subsets become much larger, those selected using imputed tracks appear to outperform those selected using the primary data. This trend may arise because imputed tracks can serve as denoised versions of the primary data [3]. At the beginning of the selection process, this denoising is not necessary to select experiments that are very different from each other. However, once many experiments have been selected, the denoised experiments may be better at identifying real differences between experiments.

**Figure 3:**
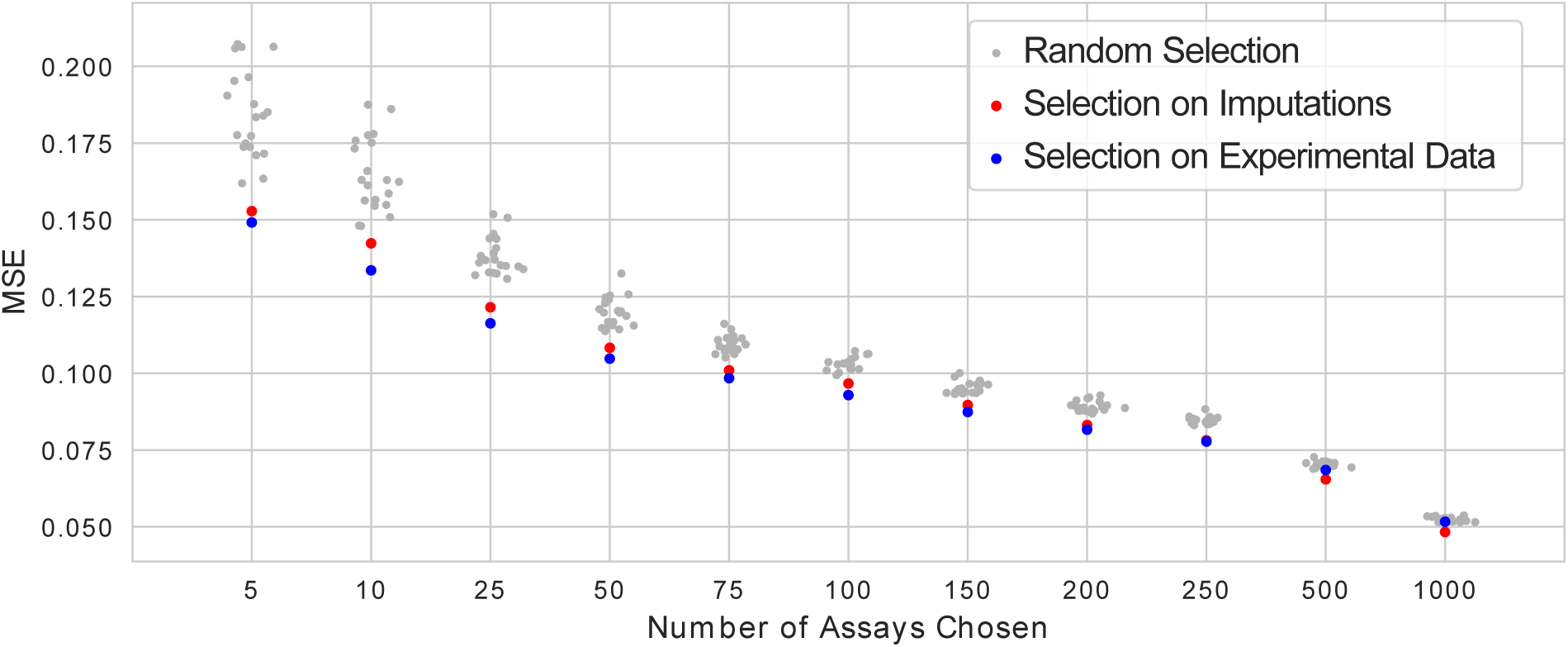
Imputation performance using different panels of assays. The performance of regression models (in terms of mean-squared-error, MSE) as a function of the number of assays chosen as the input. These panels range in size from 5 assays to 1000 assays, and are selected either randomly (grey), through a facility location function applied to imputed experiments (red), or through a facility location function applied to experimental data (blue).

### 2.4 Prioritization can be performed for individual biosamples or assays

A potential weakness in selecting experiments across both biosamples and assays is that, because the selection process is driven to select experiments with very different signal profiles, differences in the shape of the signal from each assay may dominate over differences in meaningful biology. This phenomenon is reflected in the observation that assay type is the predominant determinant of location in the UMAP embedding (Figure 1a). For example, although a difference in expression of a small number of important genes across two biosamples may be critical for certain cellular processes, the resulting assay signals are likely still more similar to each other than to an assay that measures histone modification. Thus, there may be cases where it is useful to focus prioritization efforts on a particular assay or biosample in order to factor out these effects. In our proposed approach, both can be accomplished by simply prioritizing different subsets of experiments.

First, we considered the task of prioritizing the order of biosamples in which to run a given assay. We focused on H3K27ac, a histone modification that is enriched in active enhancer elements, and chromatin accessibility as measured by DNase-seq. For both of these assays, from the full correlation matrix we extracted the submatrix of all experiments that include the assay. Reassuringly, UMAP projections of the extracted submatrices show that the experiments appear to cluster by the anatomy type of the biosample that they were performed in (Figure 4a/b). Much of this structure was not apparent in the joint projection of all experiments (Figure 1a), likely because a single visualization cannot easily capture the full complexity of such a data set. Interestingly, we note that projections share similarities across assays, such as neural and stem cell biosamples forming nearby clusters, but that there are also differences, such as heart biosamples forming a more compact cluster in the H3K27ac experiments than in the DNase experiments. When we initialize our selection procedure with the biosamples that the assays have already been performed in, we confirm that the experiments that are selected next are dissimilar to those that have already been assayed (Figure 4e/f).

**Figure 4:**
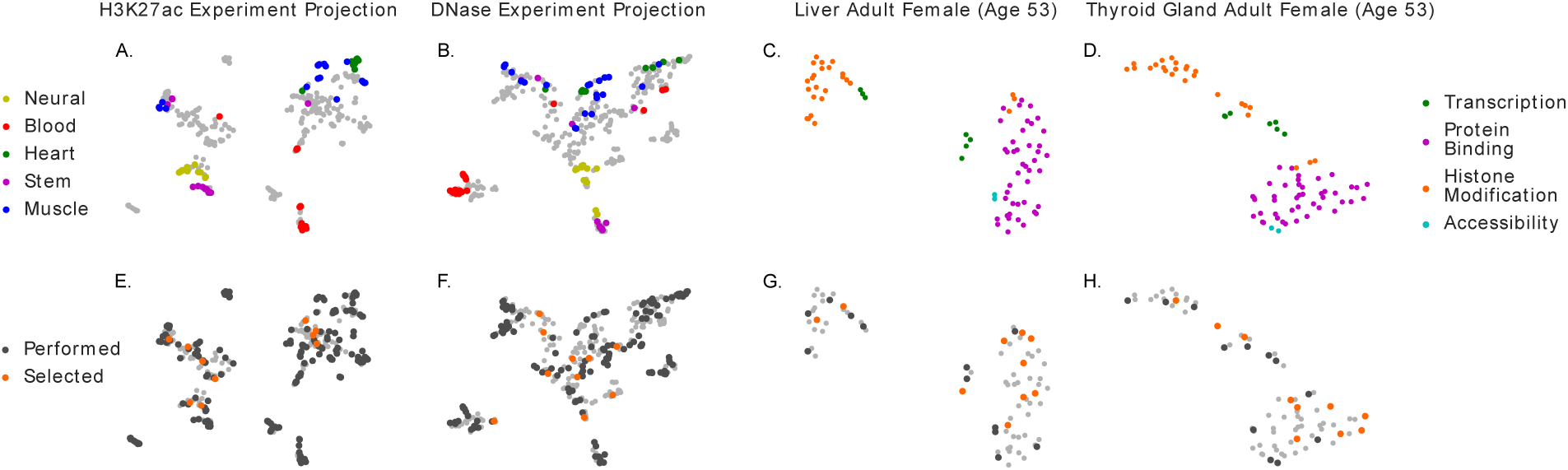
Projections and selections of all experiments containing a specific biosample or assay. UMAP projections for sets of experiments that each contain a particular biosample or assay. (A) A projection of H3K27ac experiments in all 400 biosamples, with some experiments colored according to anatomy type. (B) Same as (A), but with DNase experiments. (C) A projection of all assays performed in liver biosample, with assays colored by activity type. (D) Same as (C), but in a thyroid gland biosample from the same individual. (E-H) The same projections as (A-D), but performed experiments are colored in dark grey, the next 10 selected experiments are colored in orange, and experiments that are not selected and have not yet been performed are colored in light grey.

Next, we can also prioritize the order that assays should be performed in a given biosample by using the submatrix of experiments that include the relevant biosample. In order to highlight that Kiwano accounts for the content of the performed experiments rather than just those that have been performed, we selected two biosamples from the same individual which had the same set of experiments performed. A projection of their respective experiments resembles our joint visualization of all experiments in some aspects (Figures 4c/d). Specifically, accessibility and protein binding assays appear to form one large cluster, histone modification experiments form another cluster, and transcription assays are separate from the others but do not form their own cluster.

Similar to the joint selection procedure, we can account for experiments that have already been performed. When we prioritize the order that biosamples should be assayed, we observe that the experiments that are selected appear to be dissimilar than those that have already been assayed, and that well characterized areas are not selected from (Figure 4e/f). Likewise, despite the same set of assays having been performed in the two biosamples we considered, we observe that our procedure selects different assays (Figure 4g/h).

### 2.5 Calculating the coverage of each biosample and assay

Thus far we have focused our efforts on prioritizing individual experiments but have provided little guidance for how to prioritize entire biosamples or assays. We next considered a scenario where an investigator is looking to either assay undercharacterized biosamples or to run underperformed assays, but is unsure which biosamples or assays to focus on. A simple approach would be to count the number of experiments that each biosample or assay is involved in and choose the ones with the fewest experiments. However, this approach does not account for the content of the performed experiments, which can be extremely similar in some cases. For example, in the ENCODE data several biosamples have been assayed extensively for transcription but not assayed at all for histone modifications or protein binding.

A final component of our methodology is the ability to quantify the extent to which each biosample has been characterized and each assay has been performed using the facility location objective function. Because the objective function takes in a set of experiments and returns a score corresponding to the diversity of the set, this function can be used to assess the diversity obtained by an existing set of experiments, corresponding to a single biosample or a single type of assay. In our setting, where similarity is measured via squared correlation, this score ranges from zero up to the total number of experiments that have been performed. Thus, for each biosample, the maximum value is 77 due to the 77 assays in the data set, and for each assay, the maximum value is 400 due to the 400 biosamples in the data set.

We applied this approach to score each of the biosamples and assays in the ENCODE2018-Core data set. Not surprisingly, we find that the three ENCODE Tier 1 cell lines—H1-hESC, K562, and GM12878—are the three best scoring biosamples, with scores of 71.2, 70.2, and 68.6, respectively. These biosamples are followed by several ENCODE Tier 2 cell lines, such as HepG2, IMR90, and HeLa-S3. We found a rank correlation of 0.82 between the number of assays performed in a biosample and the objective score, confirming that while in general there is increase in coverage as more assays are performed, the composition of those assays is also captured by the objective function. Next, we scored the assays and found that the highest scoring ones were H3K4me3, H3K36me3, and CTCF, whereas the lowest scoring assays are H2BK15ac and FOXK2. We found a weaker, but still very significant, rank correlation of 0.66 between the number of biosamples that an assay was performed in and the objective score.

We next sought to contextualize the scores we obtained for each biosample and assay by comparing them to scores obtained if one had used alternate methods to select experiments. For each element, i.e., a particular assay or biosample, we scored 10 randomly selected panels of the same size as the number of experiments involving that element. Additionally, we score the panel of experiments that would have been selected using submodular selection. We observe a striking result, which is that the set of experiments that were actually performed not only underperforms the set selected through submodular selection, but also generally underperform random selection (Figure 5). This trend is consistent across both biosamples and assays. We note that the 64 biosamples with the worst scores were assayed almost exclusively for transcription, supporting the notion that biosamples with more assays performed in them are not always better characterized.

**Figure 5:**
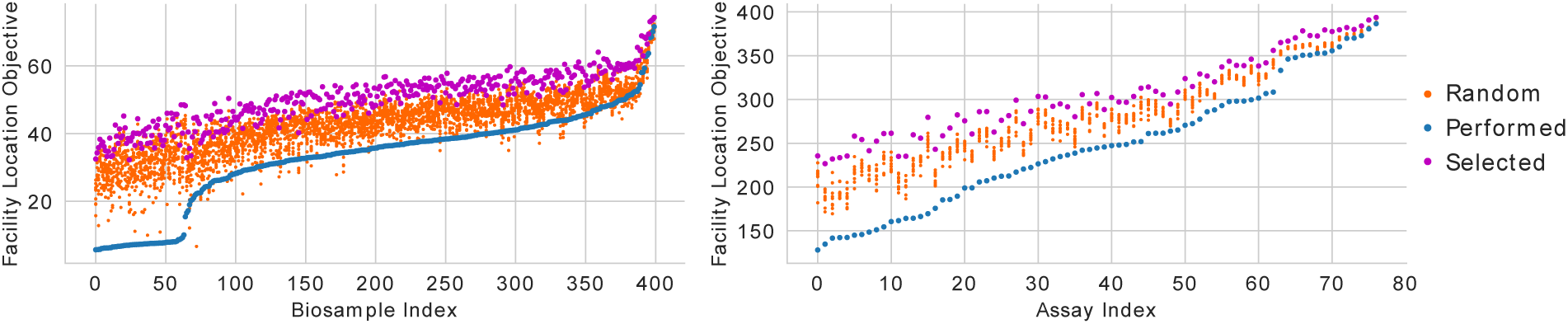
Scoring biosamples and assays according to their captured diversity. (A) The facility location objective score for each biosample when applied to the set of experiments that investigators have performed in that biosample (blue), the set of experiments identified by optimizing the objective function (magenta), and the sets of randomly selected experiments (orange), ordered by the score of the performed experiments. (B) The same as (A), but for each assay instead of each biosample.

## 3 Discussion

In this work, we describe an approach for the prioritization of epigenomic and transcriptomic experiments that has the potential to increase the rate of scientific discovery by focusing characterization efforts on those experiments that are expected to yield the least redundant information. To our knowledge, this is the first approach that enables the global prioritization of experiments across both biosamples and assays. We anticipate that, due to the time it takes to perform experiments and the simplicity of Kiwano, investigators may use our prioritization methods even when they plan eventually to perform all potential experiments in order to begin analyses sooner.

An important consideration is that, due to the reliance on imputed experiments, Kiwano cannot be applied directly to a biosample or assay type when no experiments have yet been performed. However, because a diversity of biosamples have already been experimentally characterized, in many cases it would be simple to identify closely related experiments that imputations have already been generated for. While these imputations may not capture activity specific to an experiment, it is likely that the resulting similarity matrix is still a reasonable approximation. Unfortunately, similar imputations are unlikely to be readily available in cases where one is performing experiments that are very unlike anything that have been performed before. In this setting, it would likely be necessary to first perform a subset of experiments that include all assays and biosamples for use in training an imputation model, and then use the resulting imputations to prioritize the remaining experiments.

While the primary question that we address is how to prioritize experiments across both biosamples and assays, we recognize that this approach may not always result in a practical set of experiments to perform. Specifically, it is generally more difficult to culture and maintain a variety of biosamples than it is to maintain a large quantity of a single biosample, making sets of experiments that span several biosamples harder to perform than those in the same biosample. This difficulty may cause investigators to prefer performing batches of experiments within a biosample, and so we have provided methods both for choosing biosamples that currently are not well characterized and for prioritizing assays within a given biosample. Fortunately, such difficulties are significantly reduced when performing different types of ChIP-seq assays in the same biosample, due to their standardized nature.

When we scored the biosamples that are part of the ENCODE2018-Core data set using the facility location objective function, we noted that the set of assays performed in many biosamples performed worse than randomly selecting an equally sized panel of assays. However, this trend is not entirely surprising. The experiments that are included in our data set were devised by scientists to investigate specific research questions, and generally these questions do not aim to broadly characterize the human epigenome. Thus, these results serve primarily to demonstrate that the current strategy for selecting experiments is not well suited for the goal of characterizing the overall diversity of activity in the human epigenome, and the scores can serve as a basis for prioritizing entire biosamples or assays.

A weakness in Kiwano is that mistakes in the imputation process are propagated to the selection process. These mistakes can be simple errors in predicting certain peaks, but can also involve more systematic trends. For example, REST is a transcription factor that is involved in suppressing neuronal genes in non-neuronal tissues. However, the ENCODE2018-Core data set does not have examples of REST in neuronal tissue, and so an imputation model trained on this data set would likely be unaware of this property of REST. Consequently, the prioritization process is unlikely to capture that REST binding in neuronal tissues is significantly different than in non-neuronal tissues. In general, unexpected patterns in data that has not yet been collected will be difficult for any prioritization method to account for.

The flexibility of Kiwano allows for several extensions that we did not consider here, but may nonetheless prove valuable to those prioritizing experiments. The first is that, in the setting where one is prioritizing experiments within a particular biosample, one could measure the gain that each successive experiment adds to the objective function to determine when to stop performing experiments. This would serve as a data-driven indicator of when further experimental efforts are mostly redundant. A second potential extension is to add regularization to the selection process itself to encourage successive experiments to come from the same biosample. While there are many ways that one could do this, a simple approach would be to rephrase the objective function as *f* (*X*) + *λG*(*X*), where *f* (*X*) is the facility location function as used here, *λ* is the regularization strength, and *G*(*X*) is a submodular function counting the number of biosamples not considered by this set. Because the sum of two submodular functions is itself submodular, a similar greedy optimization approach could be applied here. A third extension is that one could calculate the similarity matrix using only a specific genomic locus or set of loci of interest. For example, if an investigator was aiming to experimentally quantify the activity surrounding an important gene across many biosamples, one could restrict the similarity calculation to a window surrounding that gene. Overall, Kiwano is a simple yet powerful way to prioritize experiments in a wide variety of contexts.

## 4 Methods

### 4.1 Datasets

We generated our imputations using an Avocado model that had previously been trained on the ENCODE2018-Core data set [6]. The model is available at https://noble.gs.washington.edu/~jmschr/mango/models/. This model was trained on 3,814 experiments across 400 biosamples and 84 assays where the signal was −log_10_ p-values that had subsequently been arcsinh transformed to reduce the effect of outliers. The resulting imputations are in the same space. Due to the large size of the genome, we only imputed the ENCODE Pilot Regions, comprising ∼ 1% of the genome [11], for each of the 33,600 potential experiments.

An important detail is that, at the time of accession, experiments meausuring transcription had been divided into plus-strand signal and minus-strand signal on the ENCODE portal. Consequently, each strand was counted as a separate assay when training the Avocado model. While the strand that transcription occurs on is important for an imputation approach to capture, this distinction is not helpful for prioritizing experiments because one generally cannot perform an experiment measuring transcription on only one of the strands. Thus, we combine the plus- and minus-strand experiments for both the imputed and the primary epigenomic data by simply adding the tracks together. This process reduced the total number of assays from 84 to 77, the total number of performed experiments from 3,814 to 3,510, and the total number of potential experiments from 33,600 to 30,800.

### 4.2 Facility location and submodular optimization

Submodular optimization is the discrete analog of convex optimization and operates on submodular set functions. A function is submodular if and only if it has the property of diminishing returns; i.e., the incremental gain in function value associated with adding an element s to a set A becomes smaller as the size of the set A becomes larger. More formally, given a finite set *S* = {*s*_1_, *s*_2_, …, *s*_*n*_}, a discrete set function *f* : 2*^S^* → ℝ is submodular if and only if

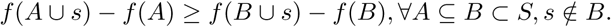

In this work, we employ a submodular function whose value is inversely related to the redundancy within a given set. Thus, optimizing such a function, subject to a cardinality constraint, involves identifying the subset whose elements are minimally redundant with each other. For further reading on submodular optimization, we suggest [12, 13, 14].

Kiwano relies on optimizing a particular submodular function called facility location. Facility location takes the form

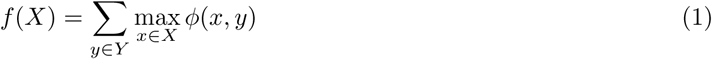

such that *Y* is the full set of experiments, *X* is the selected subset of experiments such that *X* ⊆ *Y*, *x* and *y* are individual experiments in *X* and *Y* respectively, and *ϕ*(*x, y*) is the squared correlation between *x* and *y*. The facility location function is optimized using the accelerated (or “lazy”) greedy algorithm [15]. We use apricot v0.3.0 to perform this selection [16].

### 4.3 Model training

The machine learning models used for evaluation of the selection procedure are implemented using keras (v2.2.4) [17] with a Theano (v1.0.4) [18] backend. Each model is a multi-task linear regression model, meaning that the model contains a single weight matrix that transforms inputs into outputs. Training is performed using the Adam optimizer [19] with a mean-squared-error loss for predicting epigenomic signal and a binary cross-entropy loss for predicting enhancer activity. The optimizer hyperparameters and the weight initializations are set to the keras defaults. No explicit regularization is used.

